# Interpretable factor models of single-cell RNA-seq via variational autoencoders

**DOI:** 10.1101/737601

**Authors:** Valentine Svensson, Lior Pachter

## Abstract

Single cell RNA-seq makes possible the investigation of variability in gene expression among cells, and dependence of variation on cell type. Statistical inference methods for such analyses must be scalable, and ideally interpretable. We present an approach based on a modification of a recently published highly scalable variational autoencoder framework that provides interpretability without sacrificing much accuracy. We demonstrate that our approach enables identification of gene programs in massive datasets. Our strategy, namely the learning of factor models with the auto-encoding variational Bayes framework, is not domain specific and may be of interest for other applications.

## Introduction

The study of the regulatory architecture of cells has revealed numerous examples of co-regulation of transcription of large numbers of genes (Jang *et al*., 2017; Kondo *et al*., 2018), and this has been used to link the organization of cells to their distinct functions in response to developmental or external stimuli (Romero *et al*., 2012). While studies of cells in bulk have led to interesting population-level insights about the relationships between genes (Thompson *et al*., 2015), the study of individual cells via single-cell RNA-seq has led to questions about how the relationships between genes depend on cell type (Lindgren *et al*., 2017).

Principal component analysis (PCA) is a popular linear method for dimensionality reduction in single-cell RNA-seq (Rostom *et al*., 2017; Andrews and Hemberg, 2017). As a result of its efficiency, PCA has been used for exploratory data analysis to quickly visualize the structure of high-dimensional data in two or three dimensions. PCA also provides a linear model of the data; a key feature of the method that can be used for prediction (Tipping and Bishop, 1999). In the case of single-cell RNA-seq, data points correspond to cells and the coordinates of each cell represent the gene expression levels for each gene in the transcriptome. Thus, PCA can be used to study structured variation between cells by revealing differences along axes of greatest variation. In PCA, linear weight parameters (loadings) are used to predict gene expression in each cell, conditional on the latent variables (coordinates) per cell. The loadings corresponding to the principal component axes can be interpreted as “meta-genes”: sets of genes which tend to be expressed together (Brunet *et al*., 2004; Raychaudhuri *et al*., 2000). Thus, principal component analysis of gene expression provides a formal mathematical framework for studying the biological idea of “gene programs” (Stuart *et al*., 2003) by simultaneously explaining structured variation between cells and between genes (Islam *et al*., 2011; Guo *et al*., 2010).

While PCA is easy to use and is often applied to single-cell RNA-seq data, the method has some drawbacks. PCA models data as arising from a continuous multivariate Gaussian distribution, and thus optimizes a Gaussian likelihood (Pearson, 1901; Tipping and Bishop, 1999). This model assumption is at odds with the count data measured in single cell RNA-seq (Svensson; William Townes *et al*., 2019), and leads to interpretation problems (Hicks *et al*., 2018). To address this issue, a number of methods define factor methods tailored to single cell transcriptomics data (Pierson and Yau, 2015; Zhu *et al*., 2017; Durif *et al*., 2019). For example ZINB-WaVE defines a linear factor model where gene weights are parameters, cell factor values are latent variables, and data arises from a zero-inflated negative binomial distribution (Risso *et al*., 2018). However, as single cell transcriptomics datasets have grown in size to hundreds of thousands of observations (Svensson *et al*., 2018), efficiency and scalability considerations have become paramount and inference with parametric models can be intractable. To address scalability requirements, new methods based on variational autoencoders have been developed; these leverage the large amounts of available data to learn nonlinear maps, and crucially scale well thanks to efficient algorithms for inference that leverage the structure of autoencoders (Lopez *et al*., 2018; Eraslan *et al*., 2019).

Autoencoders consists of a pair of functions: a representation function and a reconstruction function, which are typically parameterized as neural networks (Hinton and Zemel, 1994). Autoencoders can be seen as a nonlinear generalization of PCA, which can be viewed of consisting of two projections (Plaut, 2018). By optimizing the pair of neural networks, efficient low-dimensional representations of data can be identified. A variational autoencoder (VAE) uses a similar strategy but with latent variable models (Kingma and Welling, 2013). Each datapoint is represented by a set of latent variables, which can be decoded by neural networks to produce parameters for a probability distribution, forming a generative model. To infer the latent variable values (the representation), a neural network is used to find per-datapoint parameters for a probability distribution in the representation space. This defines an “inference model” which attempts to approximate the posterior distribution of the latent variables given the observed data with a variational distribution (Marino *et al*., 2018).

Inference using VAEs scales to arbitrarily large data since mini-batches of data can be used to train the parameters for both the inference model and the decoder function (Kingma and Welling, 2013). We show that using a flexible non-linear inference model along with a linear reconstruction function makes it possible to benefit from the efficiency of VAEs, while retaining the interpretability provided by factor models. Specifically, by adapting the method of scVI (Lopez *et al*., 2018), we demonstrate a scalable approach to learning a latent representation of single-cell RNA-seq data, that identifies the relationship between cell representation coordinates and gene weights via a factor model. Our approach results in a tradeoff: whereas typically autoencoder models are designed with the same network topology in the inference functions and the reconstruction functions, what we propose is a less flexible reconstruction function that suffers an increase in reconstruction error, yet provides an interpretable link between gene programs and cellular molecular phenotypes **(Figure 1a)**.

**Figure 1.**
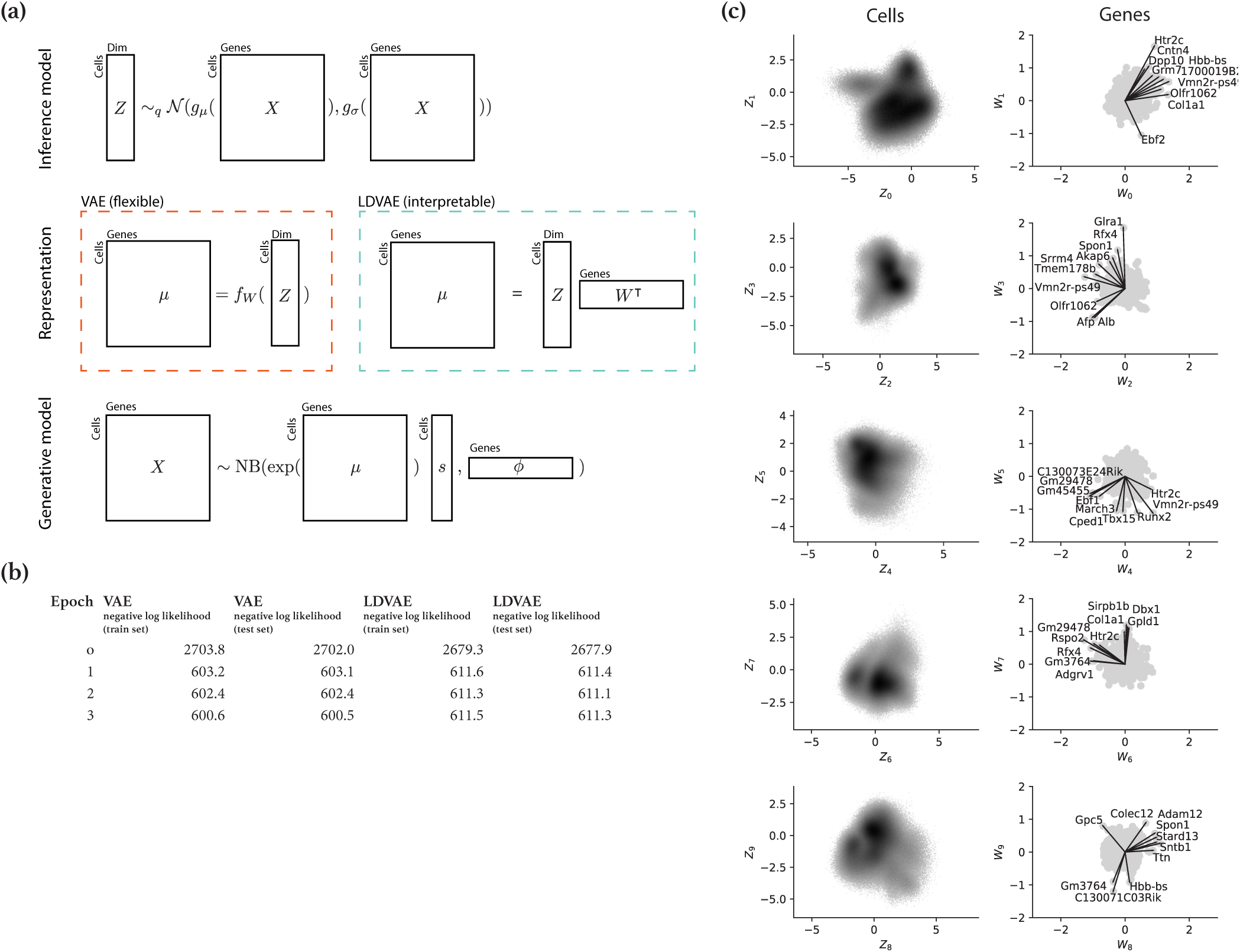
Linearly decoded variational autoencoder model. **(a)** A sketch of the general architecture of scVI autoencoders with two alternative representation models. **(b)** Comparison of reconstruction error during inference on Cao data with VAE and LDVAE. **(c)** Results from fitting a 10-dimensional LDVAE. (left column): Density plots of the 2 million cells in representation space. (right column): Scatter plots of gene loadings corresponding to the representation coordinates. Top genes indicated as vectors with names.

The generative model in scVI, when data is from a single batch and zero-inflation is deactivated, is

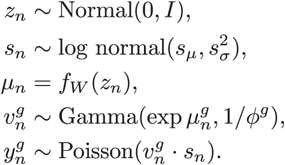

In this model *s*_*n*_ is a random variable for the exposure or count depth of a cell, with priors *s*_*µ*_ and *s*_σ_. The random variable *z*_*n*_ provides a *D*-dimensional representation of cells. The parameter *ϕ*^*g*^ represents the overdispersion of a gene. Our proposial is to replace the neural network with a linear function:

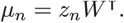

This way the expression level 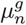 of a gene *g* in cell *n* is affected by the weights 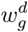depending on the coordinate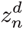 of a cell *n*, giving a direct link between cell representation and gene expression.

## Results

To investigate the potential for interpretability in the VAE framework, we implemented a linearly decoded variational autoencoder (LDVAE) in scVI. The model was applied to a recent dataset of single-cell RNA-sequencing from a large number of developing mouse embryos in different stages of development (Cao *et al*., 2019). The dataset consists of 2 million cells from 61 embryos in total, and is to our knowledge the largest scRNA-seq study published to date. After running the inference for three epochs, we confirmed that the LDVAE does indeed have a larger reconstruction error than the standard VAE **(Figure 1b)**. However, the axes of representation learned by the LDVAE model can be directly related to axes of co-expressed genes **(Figure 1c)**. For example, variation along the axis is related to simultaneous variation in expression of Tbx15 and Runx2, two genes important for cranial skeleton development (Choi *et al*., 2005; Singh *et al*., 2005). As another example variation along the axis is related to co-variation in alpha-fetoprotein (Afp) and albumin (Alb) which are related to liver development (Nayak and Mital, 1977).

To investigate the scalability, cells were subsampled to different numbers before fitting LDVAE. We found that inference runs in linear time, with 5 seconds per 1,000 cells to reach 10 epochs using a CPU (Intel Core i7-7800X). Using a consumer grade GPU (NVIDIA GeForce RTX 2070), inference only requires 2 seconds per 1,000 cells to reach 10 epochs, with a total time of less than an hour for the full dataset **(Figure 2a)**. Investigating the reconstruction error curves per epoch, the models had converged after 2-3 epochs for datasets larger than 100,000 cells **(Figure 2b)**. Determining a minimal number of epochs is a difficult general problem, but our results suggest a rule of thumb of “1 million divided by the number of cells in the dataset” epochs for first pass analysis.

**Figure 2.**
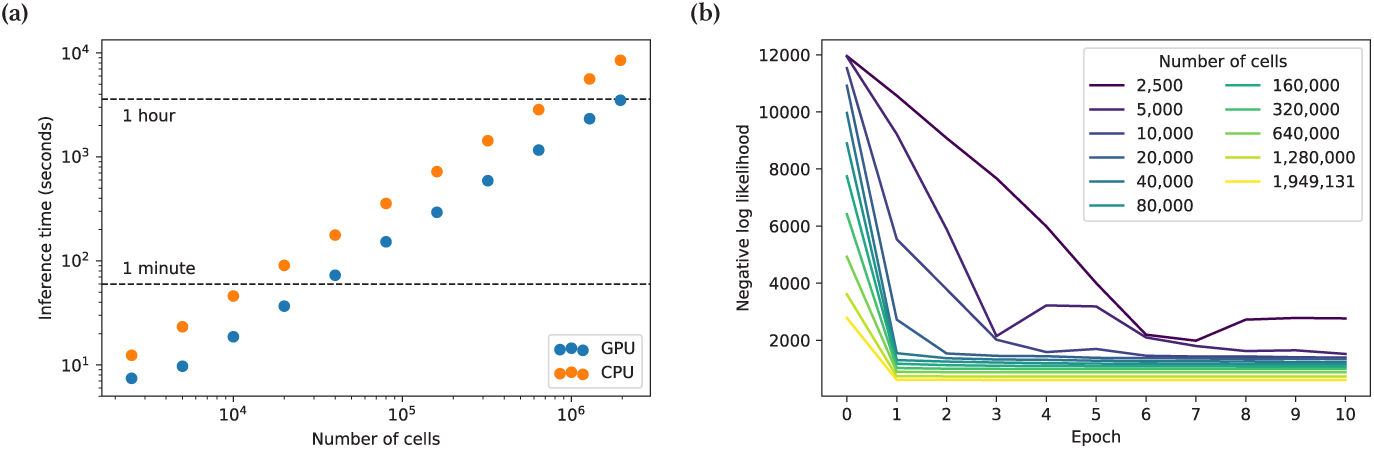
Inference speed for LDVAE. **(a)** Runtimes to reach 10 epochs with and without GPU for increasing numbers of cells. **(b)** Comparison of reconstruction error per epoch during inference for datasets with different numbers of cells.

Jupyter notebooks to produce the results are available at https://github.com/pachterlab/SP_2019 as well as CaltechDATA at http://dx.doi.org/10.22002/D1.1266. For convenience, the embryo data from Cao *et al*. are also available in an H5AD object on the CaltechDATA accession and on Google Cloud Storage at gs://h5ad/2019-02-Cao-et-al-Nature/cao_atlas.h5ad. A general tutorial on how to use the LDVAE model is available in the scVI GitHub repository at https://github.com/YosefLab/scVI/blob/master/tests/notebooks/linear_decoder.ipynb.

## Discussion

Our results show that interpretable non-Gaussian factor models can be linked to variational autoencoders to enable interpretable analysis of data at massive scale. This is useful for the investigation of gene co-expression in large scRNA-seq datasets, and the approach we have outlined should be applicable in other settings where interpretability is paramount. The development of our approach utilized a well-documented implementation of VAEs for single-cell RNA-seq in the scVI package (Lopez *et al*., 2018), and the development of the LDVAE model was greatly facilitated by the ability to build on the existing codebase of scVI; the methods we have described have been merged into the scVI code base. We believe that our work provides a useful example of the value of building on existing frameworks in bioinformatics, and specifically, in the case of single-cell RNA-seq, we believe that other extensions of the scVI VAE models could be similarly fruitful to explore.

## Acknowledgements

We thank Eduardo da Veiga Beltram and Romain Lopez for helpful feedback on the manuscript. Additionally we thank the developers and users of scVI who provided helpful discussion and feedback for the implementation on Github. The work was partly funded by NIH U19MH114830.

## Notes

http://dx.doi.org/10.22002/D1.1266

https://github.com/pachterlab/SP_2019

